# Physical activity and risk of lung cancer: a two-sample Mendelian randomization study

**DOI:** 10.1101/806083

**Authors:** Sebastian E Baumeister, Michael F Leitzmann, Martin Bahls, Christa Meisinger, Christopher I Amos, Rayjean J Hung, Cancer in Lung of the International Lung Cancer Consortium (TRICL-ILCCO), Lung Cancer Cohort Consortium (LC3), Alexander Teumer, Hansjörg Baurecht

**Author notes:** contributed equally (Alexander Teumer, Hansjörg Baurecht). Collaborators of the TRICL-ILCCO and LC3 consortium are listed in this article’s supplementary note. **Corresponding author:** Sebastian E. Baumeister, PhD, Chair of Epidemiology, Ludwig-Maximilians-Universität München, UNIKA-T Augsburg, Neusässer Str. 47, 86156 Augsburg, Germany.

## Abstract

Observational studies have suggested that physical activity might lower the risk of lung cancer in former and current smokers but not in never smokers. Using genetic instruments for self-reported and accelerometer-measured physical activity traits implemented through two-sample Mendelian randomization (MR), we sought to strengthen the evidence for causality. We used 18 genome-wide significant (P < 5×10^−8^) single nucleotide polymorphisms (SNPs) for self-reported moderate-to-vigorous physical activity and seven SNPs for accelerometer-measured (‘average acceleration’) physical activity from up to 377,234 UK Biobank participants and evaluated these in relation to risk using 29,266 lung cancer cases (including 11,273 adenocarcinomas, 7,426 squamous cell and 2,664 small cell cases) and 56,450 controls. The MR analysis suggested no effect of self-reported physical activity (odds ratio (OR) [95% confidence interval (CI)] = 0.67 [0.42-1.05], P-value = 0.081, Q-value = 0.243) and accelerometer-measured activity (OR [95% CI] = 0.98 [0.93-1.03], P-value = 0.372,Q-value = 0.562) on lung cancer. There was no evidence for associations of physical activity with histologic types and lung cancer in ever and never smokers. Replication analysis using genetic instruments from a different genome-wide study and sensitivity analysis to address potential pleiotropic effects led to no substantive change in estimates. These findings do not support a protective relationship between physical activity and the risk of lung cancer.

**Significance:** The present study provides little evidence that recommending physical activity would help to prevent lung cancer.

## Introduction

Lung cancer is the leading cause of cancer mortality worldwide (1). Although smoking is the risk factor most strongly linked to all lung cancer subtypes, about 10% of cases are seen in never-smokers (2). Potential non-smoking related risk factors for lung cancer include environmental carcinogens, pulmonary fibrosis, genetic history, dietary factors, and insufficient physical activity (3,4). Several meta-analyses of observational studies suggested an inverse association between physical activity and lung cancer risk (5-7). Yet, the evidence has been limited to current and former smokers in most studies (5-7). Interpretation of this inverse association has been constrained by potential confounding, as smoking causes lung cancer and renders physical activity more difficult (5,8). Reverse causation may also affect the association between physical activity and lung cancer risk, as the presence of lung cancer symptoms may lead to avoidance of physical activity (9). Accordingly, the World Cancer Research Fund/American Institute for Cancer Research (4) and a recent umbrella review (10) have categorized the overall evidence from observational studies as inconclusive. Mendelian randomization (MR) is a method that uses genetic variants as instrumental variables to help uncover causal relationships in the presence of unobserved confounding and reverse causation (11). In the current study, we performed two-sample summary data MR analyses to assess the association between physical activity and lung cancer.

## Methods

The study had five components: (1) identification of genetic variants to serve as instrumental variables for physical activity traits; (2) acquisition of instrumenting SNP-outcome summary data from genome wide association studies (GWAS) of lung cancer; (3) harmonization of SNP-exposure and SNP-outcome datasets; (4) statistical analysis; (5) evaluation of MR analysis assumptions and sensitivity analyses.

### Physical activity measurement in UK Biobank

Data for the genetic associations with self-reported and accelerometer-based physical activity phenotypes were obtained from two published GWAS conducted in the UK Biobank (12,13). The UK Biobank study is a community-based prospective cohort study that recruited over 500,000 men and women aged 40-69 years from different socioeconomic backgrounds from 22 centers across the United Kingdom between 2006 and 2010 (14). For the first GWAS by Klimentidis et al. (13), self-reported levels of physical activity were ascertained in 377,234 UK Biobank participants using the International Physical Activity short form Questionnaire (15) and moderate-to-vigorous physical activity was computed by taking the sum of total minutes per week of moderate and vigorous physical activity multiplied by eight, corresponding to their metabolic equivalents (13). For objective assessment of physical activity, a subset of 103,712 participants wore an Axivity AX3 triaxial accelerometer on the wrist for a seven-day-period between 2013 and 2015 (16). After calibration, removal of gravity and sensor noise, and identification of wear/non-wear episodes the remaining 100Hz raw triaxial acceleration data was used to calculate physical activity variables. Non-wear time was defined as consecutive stationary episodes lasting for at least 60 minutes where all three axes had a standard deviation of less than 13.0 milli-gravities. For the GWAS by Klimentidis et al. (13), ‘average acceleration’ (in milli-gravities) was used as the exposure variable derived from accelerometer wear. For the second GWAS by Doherty et al. (12), accelerometer-measured ‘overall activity’ levels were defined as average vector magnitude for each 30-s epoch (16). Written informed consent was obtained from UK Biobank study participants and ethics approval of UK Biobank was given by the North West Multicentre Research Ethics Committee, the National Information Governance Board for Health & Social Care and the Community Health Index Advisory Group. Both GWAS studies (12,13) were covered by the general ethical approval of the UK Biobank studies from the NHS National Research Ethics Service on 17th June 2011 (Ref 11/NW/0382).

### Selection of genetic instrumental variables for physical activity

For the primary analysis, we initially selected 19 SNPs associated with self-reported moderate-to-vigorous physical activity at a genome-wide significance level (P < 5 × 10^−8^) in the GWAS by Klimentidis et al. (13), using The PLINK clumping algorithm (r^2^ threshold = 0.001 and window size = 10mB) (Supplementary Table 1). We identified eight SNPs associated with accelerometer-measured ‘average acceleration’ at P < 5 × 10^−8^(13) (Supplementary Table 2). For the secondary analysis, we selected six SNPs associated with accelerometer-measured ‘overall activity’ at < 5 × 10^−8^ in the GWAS by Doherty et al. (12) (Supplementary Table 2). After removal of SNPs exhibiting potential pleiotropic effects (see details in ‘Statistical analyses’ and ‘Results’), 18, 7 and 5 SNPs were used as instruments for self-reported moderate-to-vigorous physical activity, accelerometer-measured ‘average acceleration’ and accelerometer-measured ‘overall activity’, respectively. UK Biobank participants were genotyped using the UK BiLEVE array and the UK Biobank axiom array. Tables 1 and 2 present the harmonized genetic instruments and associations with physical activity traits.

**Table 1.**
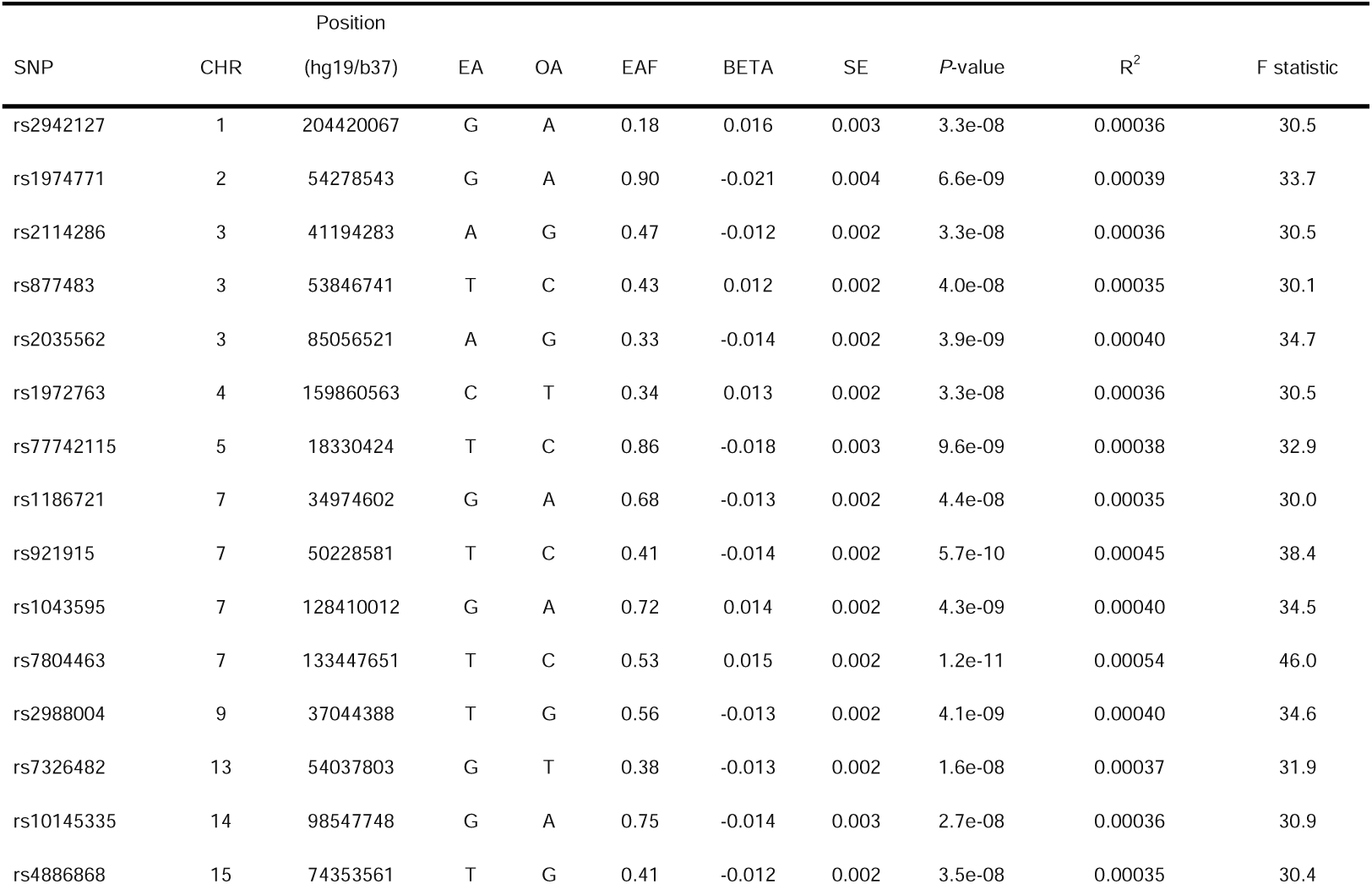
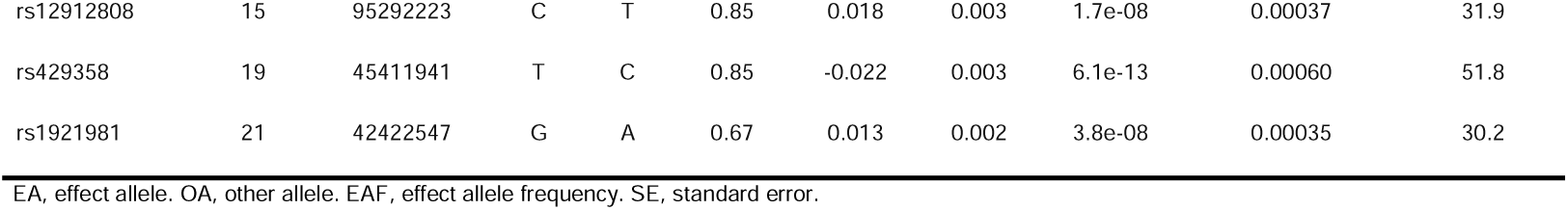
Self-reported moderate-to-vigorous physical activity SNPs from the GWAS by Klimentidis et al. (13) used as genetic instruments in the primary Mendelian analysis

**Table 2.** Accelerometer-measured physical activity SNPs used as genetic instruments in the primary and secondary Mendelian analysis

| SNP | CHR | Position | EA | OA | EAF | BETA | SE | P-value | R<sup>2</sup> | F |
| --- | --- | --- | --- | --- | --- | --- | --- | --- | --- | --- |
|  |  | (hg19/b37) |  |  |  |  |  |  |  |  |
| Primary analysis (SNPs from GWAS by Klimentidits et al. (13) – ‘average acceleration’) |  |  |  |  |  |  |  |  |  |  |
| rs336605 | 3 | 18656350 | G | T | 0.28 | 0.222 | 0.041 | 4.5e-08 | 0.00035 | 29.9 |
| rs10067451 | 5 | 87942506 | G | A | 0.89 | 0.326 | 0.058 | 2.5e-08 | 0.00036 | 31.1 |
| rs28749810 | 5 | 152048630 | C | A | 0.66 | 0.210 | 0.038 | 4.4e-08 | 0.00035 | 30.0 |
| rs7084454 | 10 | 21821274 | G | A | 0.68 | 0.222 | 0.039 | 1.0e-08 | 0.00038 | 32.8 |
| rs148193266 | 11 | 104528681 | A | C | 0.96 | -0.510 | 0.092 | 3.1e-08 | 0.00036 | 30.7 |
| rs79724577 | 17 | 43463493 | A | C | 0.82 | -0.276 | 0.047 | 4.6e-09 | 0.00040 | 34.4 |
| rs1518139 | 18 | 40751232 | G | T | 0.66 | -0.226 | 0.039 | 4.5e-09 | 0.00040 | 34.4 |
| Secondary analysis (SNPs from GWAS by Doherty et al. (12) – ‘overall activity’) |  |  |  |  |  |  |  |  |  |  |
| rs6775319 | 3 | 18758501 | A | T | 0.27 | 0.027 | 0.005 | 3.9e-08 | 0.00035 | 30.2 |
| rs9293503 | 5 | 87948962 | T | C | 0.89 | 0.039 | 0.007 | 4.9e-08 | 0.00035 | 29.8 |
| rs6873698 | 5 | 152039420 | C | T | 0.66 | 0.027 | 0.005 | 2.6e-08 | 0.00036 | 31.0 |
| rs11012732 | 10 | 21830104 | A | G | 0.67 | 0.028 | 0.005 | 4.1e-09 | 0.00040 | 34.6 |
| rs59499656 | 18 | 40768309 | A | T | 0.66 | -0.028 | 0.005 | 1.9e-09 | 0.00042 | 36.1 |

EA, effect allele. OA, other allele. EAF, effect allele frequency. SE, standard error.

### GWAS summary statistics for lung cancer

Genetic variants associated with lung cancer were obtained from a meta-analysis of GWAS (17), comprising the Lung Cancer Consortium (TRICL-ILCCO) lung cancer GWAS (11,177 lung cancer cases and 40,396 controls) (18) and an additional 18,089 lung cancers and 16,054 controls from the the Lung Cancer Cohort Consortium (LC3). The individual studies were genotyped on different arrays, imputed based on 1000 Genomes (phase 3) and harmonized (17). The overall sample size was 29,266 lung cancer cases and 56,450 controls. The GWAS analysis was stratified by histology, including 11,273 adenocarcinomas, 7,426 squamous cell carcinomas, and 2,664 small cell lung cancers. Additionally, analyses were stratified by smoking status defined as ever smoker (current and former smokers; 23,223 cases and 16,964 controls) and never smokers (2,355 cases and 7,504 controls). For all SNPs used as instruments for physical activity traits, harmonized SNP-lung cancer associations are provided in Supplementary Tables 1 and 2. The studies participating in the TRICL-ILCCO/LC3 were approved by local intern review boards or ethics commitees.

### Statistical power

The a priori statistical power was calculated according to Brion et al. (19). The self-reported moderate-to-vigorous physical activity SNPs explained 0.7% and the accelerometer-measured physical activity SNPs explained 0.3% of the phenotypic variance in the GWAS by Klimentidis et al. (13). Given a type 1 error of 5%, we had sufficient statistical power (≥80%) when the expected odds ratios (OR) per 1-SD for overall lung cancer were ≤0.80 and ≤0.68 in genetically instrumented self-reported moderate-to-vigorous physical activity and accelerometer-measured physical activity, respectively, in the primary analysis. The accelerometer-measured physical activity SNPs in the GWAS by Doherty et al. (12) explained 0.2% of the phenotypic variance and provided statistical power ≥80% (α=5%) to detect an OR per 1-SD for overall lung cancer of 0.8. Power estimates for the self-reported and accelerometer-measured physical activity by subtypes of lung cancer are presented in Supplementary Table 5.

### Statistical analyses

We adopted a two-sample summary data MR strategy to perform analysis based on GWAS summary data and used the multiplicative random effects inverse-variance weighted (IVW) and maximum likelihood methods as our principal MR analyses approaches (11,20). The IVW estimates are obtained from IVW meta-analysis of the ratio estimates from the individual variants. We conducted the multiplicative random effects IVW instead of the fixed effects IVW because it allowed for each SNP to have different mean effects (20). The multiplicative random effects model provides valid causal estimates under the assumption of balanced pleiotropy. The maximum likelihood method estimates the causal effect by direct maximization of the likelihood given the SNP-exposure and SNP-outcome effects, assuming no heterogeneity and horizontal pleiotropy. We applied the Benjamini-Hochberg procedure to adjust for multiple testing and presented Q-values (21). Results are presented as OR per 1-SD increment in self-reported moderate-to-vigorous physical activity (MET-minutes/week) or accelerometer-measured physical activity. One SD of ‘average acceleration’ in the UK Biobank Study is approximately 8 milli-gravities (or 0.08 m/s^2^) of acceleration in a mean 5-second window (13). Analyses were performed using the TwoSampleMR (version 0.5.2) (22) and MRPRESSO (version 1.0) packages in R (version 3.6.3). Reporting followed the STROBE-MR statement (23).

### Sensitivity analyses

For the estimates from two-sample MR analysis to be valid, the genetic instrumental variable must be associated with physical activity (relevance), independent of all confounders of physical activity and lung cancer (exchangeability), and independent of lung cancer given physical activity (exclusion restriction) (24). The instrument relevance was measured by calculating the F statistic (25). We checked each candidate SNP and its proxies (r^2^>0.8) in PhenoScanner (26) and the GWAS catalog (27) for previously reported associations (P<5×10^−8^) with confounders or lung cancer. We considered smoking, chronic bronchitis, tuberculosis, pulmonary function, and pneumonia as relevant confounders (3-5,28). We also performed leave-one-out analysis to assess whether the IVW estimate is driven or biased by a single SNP.

In sensitivity analyses, we conducted MR analyses robust to particular forms of potential unbalanced horizontal pleiotropy (i.e., a process by which instruments associate with other traits that influence the outcome, a form of violation of the exclusion restriction assumption) (11) using the weighted median method (11). A modified 2^nd^ order weighting approach was used to estimate the Cochran’s Q statistic as a measure of heterogeneity (29). We also assessed the presence of directional pleiotropy using MR Egger regression based on its intercept, where deviation from a zero intercept indicates pleiotropy (11). The MR-Pleiotropy RE-Sidual Sum and Outlier (MR-PRESSO) method (22,30) was used to detect and correct for outliers in the IVW linear regression.

### Data availability

The summary statistics for the physical activity GWAS by Klimentidis et al. (13) is available at https://klimentidis.lab.arizona.edu/content/data (access date: 2020/01/27) and the summary data for the GWAS by Doherty et al. (12) is available at https://doi.org/10.5287/bodleian:yJp6zZmdj (access date: 2020/03/22). The lung cancer GWAS (17) summary data is available upon request from the TRICL-ILCCO/LC3 consortium.

## Results

Self-reported physical activity was measured in 377,234 individuals in UK Biobank that had GWAS data. Accelerometer-measured physical activity was available from 91,084 individuals in UK Biobank. The mean age of study participants was 56.0 years (SD=7.9), and 54.5% were women. The mean (SD) self-reported moderate-to-vigorous physical activity was 1,650 (2,084) MET-minutes/week. The values for the accelerometer-measured physical activity exposure ‘average acceleration’ was 27.9 (27.0) milli-gravities.

### MR analysis for physical activity and lung cancer

We found that genetically predicted self-reported moderate-to-vigorous physical activity was unrelated to overall lung cancer (IVW OR per 1-SD increment: 0.67; 95% CI: 0.42-1.05, *P-*value = 0.081, *Q*-value = 0.243), to the histologic types and lung cancer in ever or never smokers (Table 3). Likewise, accelerometer-measured ‘average acceleration’ was not associated with overall lung cancer (IVW OR per 1-SD increment: 0.98; 95% CI: 0.93-1.03, *P-*value = 0.375, *Q*-value = 0.562), and in analyses by subtypes and smoking status (Table 3). In the secondary analysis, null associations for overall lung cancer, histologic types and cancer in never and ever smokers were replicated using the accelerometer-measured ‘overall accelerations’ as an exposure variable (Supplementary Tables 6).

**Table 3.** Mendelian randomization estimates for the relationship between self-reported moderate-to-vigorous physical activity and lung cancer

| Outcome | Method | OR<sup>a</sup> | (95% CI)<sup>a</sup> | P-value | Q-Value |
| --- | --- | --- | --- | --- | --- |
| Overall lung cancer | Inverse-variance weighted | 0.67 | (0.42;1.05) | 0.081 | 0.243 |
|  | Maximum likelihood | 0.67 | (0.42;1.06) | 0.090 | 0.269 |
|  | Weighted median | 0.79 | (0.39;1.58) | 0.508 | 0.610 |
|  | MR PRESSO | 0.67 | (0.42;1.05) | 0.099 | 0.610 |
| Adenocarcinoma | Inverse-variance weighted | 0.77 | (0.38;1.56) | 0.470 | 0.470 |
|  | Maximum likelihood | 0.78 | (0.41;1.48) | 0.442 | 0.442 |
|  | Weighted median | 0.58 | (0.23;1.46) | 0.250 | 0.610 |
|  | MR PRESSO | 0.77 | (0.38;1.56) | 0.480 | 0.610 |
| Squamous cell carcinoma | Inverse-variance weighted | 0.45 | (0.2;1.05) | 0.064 | 0.243 |
|  | Maximum likelihood | 0.46 | (0.22;0.97) | 0.041 | 0.245 |
|  | Weighted median | 0.44 | (0.15;1.29) | 0.134 | 0.610 |
|  | MR PRESSO | 0.45 | (0.2;1.05) | 0.081 | 0.610 |
| Small cell carcinoma | Inverse-variance weighted | 0.37 | (0.1;1.43) | 0.151 | 0.303 |
|  | Maximum likelihood | 0.38 | (0.11;1.36) | 0.137 | 0.274 |
|  | Weighted median | 0.47 | (0.08;2.87) | 0.416 | 0.610 |
|  | MR PRESSO | 0.37 | (0.1;1.43) | 0.170 | 0.610 |
| Never smoker | Inverse-variance weighted | 0.52 | (0.11;2.42) | 0.402 | 0.470 |
|  | Maximum likelihood | 0.52 | (0.12;2.25) | 0.378 | 0.442 |
|  | Weighted median | 0.44 | (0.05;3.67) | 0.447 | 0.610 |
|  | MR PRESSO | 0.52 | (0.11;2.42) | 0.414 | 0.610 |
| Ever smoker | Inverse-variance weighted | 0.73 | (0.39;1.36) | 0.320 | 0.470 |
|  | Maximum likelihood | 0.74 | (0.39;1.37) | 0.337 | 0.442 |
|  | Weighted median | 0.89 | (0.4;2) | 0.775 | 0.775 |
|  | MR PRESSO | 0.73 | (0.46;1.17) | 0.205 | 0.775 |
| MR PRESSO, MR Pleiotropy RESidual Sum and Outlier. <sup>a</sup> OR (odds ratio) per one standard deviation increment in metabolic-equivalent (MET)-minutes/week. CI, confidence interval. |  |  |  |  |  |

### Sensitivity analyses

The F statistics for all physical activity genetic instruments were 29.9 or larger consistent with an absence of weak instrument bias (Tables 1 and 2). In the PhenoScanner database, we identified one of the 19 SNPs for self-reported moderate-to-vigorous physical activity and one of the eight SNPs for accelerometer-measured ‘average acceleration’ physical activity associated with lung cancer (Supplementary Tables 3 and 4). In the secondary analysis, one of the five SNPs for accelerometer-measured ‘overall activity’ physical activity was associated with forced vital capacity (Supplementary Table 4). We removed these SNPs exhibiting pleiotropic effects from MR analyses. The effect estimates for self-reported and accelerometer-measured physical activity traits and lung cancer were similar when using methodologies that are robust to potential pleiotropy of the genetic variants used in the analysis (Tables 3 and 4). The modified Q statistic suggested no notable heterogeneity across individual SNPs (Supplementary Table 7). Furthermore, analysis leaving out each SNP and MR-PRESSO revealed that no single SNP drove the results (Tables 3 and 4, Supplementary Tables 6, 8-10). The MR Egger intercept tests suggested no directional horizontal pleiotropy (Supplementary Table 11).

**Table 4.**
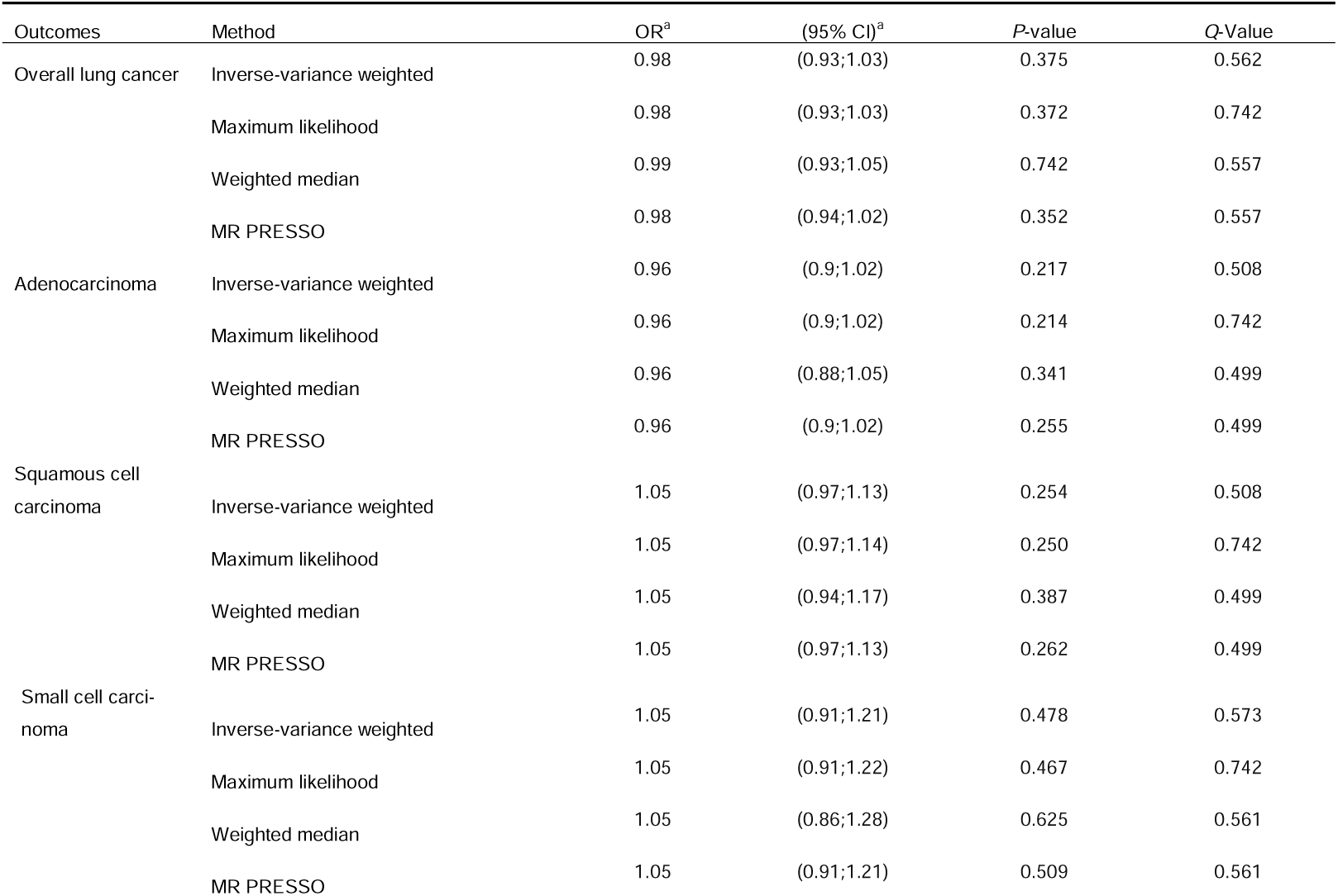
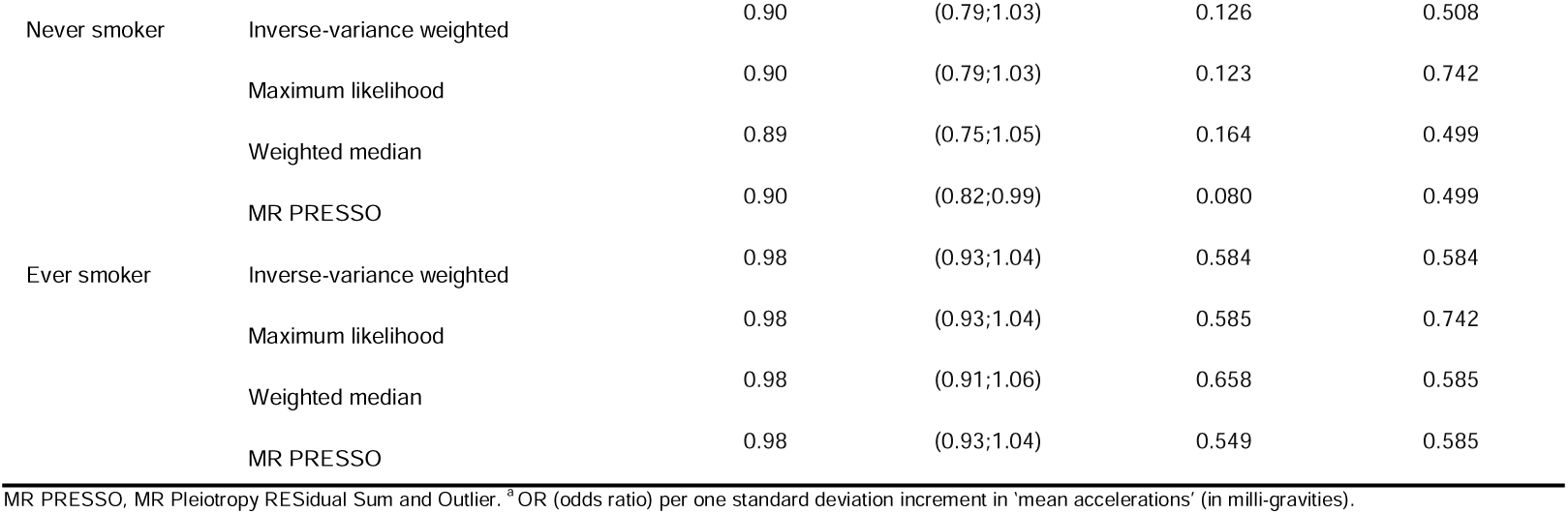
Mendelian randomization estimates for the relationship between accelerometer-measured physical activity (‘average acceleration’) and lung cancer

## Discussion

In this study, we explored the relationship of physical activity with risk of lung cancer by taking forward genetic instruments, identified in GWAS applied to approximately 377,000 UK Biobank participants, to MR analysis using data from the TRICL-ILCCO/LC3 consortium, including over 29,000 cases of lung cancer. Our principal findings suggest that physical activity (assessed using self-reported moderate-to-vigorous and accelerometer-measured activity) does not affect the risk of lung cancer. Additionally, we found no evidence for associations between physical activity and histologic subtypes and lung cancer in ever and never smokers.

In contrast to our findings, meta-analyses of observational studies concluded that higher levels of self-reported physical activity are associated with a lower risk of lung cancer (5-7). A large pooled analysis of 12 European and US cohort studies including 19,133 lung cancers reported a relative risk reduction of 24% (hazard ratio: 0.76; 95% CI: 0.71-0.77) comparing high and low levels of self-reported physical activity (31). The most comprehensive metaanalysis comprising 20 cohort studies and 31,807 cases found a 17% relative reduction in lung cancer risk with highest versus lowest levels of physical activity (hazard ratio: 0.83; 95% CI: 0.77-0.90) (7). The findings of another meta-analysis suggest no heterogeneity between histologic subtypes (5). Of note, the above-mentioned pooled analysis revealed an inverse association in current and former smokers and a null association in never-smokers (31). Similarly, meta-analyses consistently found that physical activity was inversely associated with lung cancer among former and current smokers but unrelated to lung cancer among never smokers (5-7) suggesting that negative confounding by smoking or a reduction in physical activity levels prior to diagnosis could be an explanation (8,9).

Traditional observational studies assessing the association between behavioral factors and cancers strongly associated with smoking are susceptible to confounding and reverse causation (8,32). MR offers the possibility to overcome confounding and reverse causation using genetic proxies of physical activity that are unrelated to smoking and other confounding factors when instrumental variable assumptions are fulfilled. We verified these assumptions, most notably possible pleiotropic effects, and conducted additional MR analyses using methods robust to potential unbalanced horizontal pleiotropy. The repertoire of robust MR approaches that seek to act as a sensitivity analysis (11,20,33) each makes a different series of assumptions, providing triangulating evidence (34) for our finding. The major strength of this study was the use of MR, which is less susceptible to problems of confounding, reverse causation and exposures non-differentially measured with error in comparison to conventional observational studies (35). The use of two-sample summary data MR enabled the use of the largest GWAS of lung cancer (17) to date. The study had sufficient statistical power to detect the previous observationally reported effect sizes for self-reported physical activity and overall lung cancer risk (6,7).

The study also has some limitations. First, the genetic instruments for accelerometer-assessed physical activity explained a small fraction of the phenotypic variability, which resulted in some of the subgroup analyses being underpowered. Consequently, the CIs for our MR analysis by histologic type and lung cancers in never smokers were wide. Had there been more independent genome-wide significant SNPs available that explain more of the phenotypic variability, the statistical inference could have provided more precise estimates. Second, for the two-sample MR to provide unbiased estimates, the risk factor and outcome sample should come from the same underlying population. The discovery GWAS of physical activity consisted of UK Biobank participants of European descent, aged 40 to 70 years (12,13). The SNP-lung cancer associations were derived from cohort and case-control studies of men and women of European descent aged 18 years and older (17). Given the limited age range of the UK Biobank and inclusion of European ancestry individuals only, our results may not be generalizable to other age groups or ancestral populations. Therefore, replication of our findings in other age groups and non-European populations is warranted. In conclusion, our findings provided little evidence that physical activity would help to prevent lung cancer.

## Disclaimer

The authors alone are responsible for the views expressed in this article and they do not necessarily represent the views, decisions or policies of the institutions with which they are affiliated.

## Authors’ Contributions

### Conception and design

Sebastian E Baumeister, Michael F Leitzmann, Martin Bahls, Christa Meisinger, Alexander Teumer, Hansjörg Baurecht

### Acquisition of data (provided animals, acquired and managed patients, provided facilities, etc.)

Christopher I Amos, Rayjean J Hung

### Analysis and interpretation of data (e.g., statistical analysis, biostatistics, computational analysis)

Sebastian E Baumeister, Alexander Teumer, Hansjörg Baurecht

### Writing, review, and/or revision of the manuscript

Sebastian E Baumeister, Michael F Leitzmann, Martin Bahls, Christa Meisinger, Christopher I Amos, Rayjean J Hung, Alexander Teumer, Hansjörg Baurecht

### Administrative, technical, or material support (i.e., reporting or organizing data, constructing databases)

Sebastian E Baumeister, Hansjörg Baurecht

### Study supervision

Sebastian E Baumeister, Alexander Teumer, Hansjörg Baurecht

## Acknowledgments

Transdisciplinary Research for Cancer in Lung (TRICL) of the International Lung Cancer Consortium (ILCCO) was supported by grants U19-CA148127 and CA148127S1. ILCCO data harmonization is supported by the Cancer Care Ontario Research Chair of Population Studies to R.J.H. and the Lunenfeld-Tanenbaum Research Institute, Sinai Health System. The TRICL-ILCCO OncoArray was supported by in-kind genotyping by the Centre for Inherited Disease Research (26820120008i-0-26800068-1).

