## Supplement for "Physical activity and risk of lung cancer: a two-sample Mendelian randomization study"

Sebastian E Baumeister et al.

Supplement

Cancer in Lung of the International Lung Cancer Consortium (TRICL-ILCCO) and Lung Cancer Cohort Consortium (LC3)

James D McKay^1,94^, Rayjean J Hung^2,94^, Younghun Han^3^, Xuchen Zong^2^, Robert Carreras-torres^1^, David C Christiani^4^, Neil E Caporaso^5^, Mattias Johansson^1^, Xiangjun Xiao^3^, Yafang li3, Jinyoung Byun3 , Alison Dunning^6^, Karen A Pooley^6^, David C Qian^3^, Xuemei Ji^3^, Geoffrey Liu^2^, Maria N Timofeeva^1^, Stig E Bojesen^7–9^, Xifeng Wu^10^, Loic Le Marchand^11^, Demetrios Albanes^5^, Heike Bickeböller^12^, Melinda C Aldrich^13^, William S Bush^14^, Adonina Tardon^15^, Gad Rennert^16^, M Dawn Teare^17^, John K Field^18^, Lambertus A Kiemeney^19^, Philip Lazarus^20^, Aage Haugen^21^, Stephen Lam^22^, Matthew B schabath^23^, Angeline S Andrew^24^, Hongbing Shen^25^, Yun-chul Hong^26^, Jian-min Yuan^27^, Pier Alberto Bertazzi^28,29^ , Angela c Pesatori^29^, Yuanqing Ye^10^, Nancy Diao^4^, Li Su^4^, Ruyang Zhang^4^, Yonathan Brhane^2^, Natasha Leighl^30^, Jakob S Johansen^31^, Anders Mellemgaard^31^, Walid Saliba^16^, christopher A Haiman^32^, lynne R Wilkens^11^, Ana Fernandez-Somoano^15^, Guillermo Fernandez-Tardon^15^, Henricus F m van der Heijden^19^, Jin Hee Kim^33^, Juncheng Dai^25^, Zhibin Hu25, Michael P A Davies^18^, Michael W Marcus^18^, Hans Brunnström^34^, Jonas Manjer^35^,Olle Melander^35^, David C Muller36 , Kim Overvad37, Antonia Trichopoulou^38^, Rosario Tumino39, Jennifer A Doherty^24,40–42^, Matt P Barnett^40^, Chu Chen^40^, Gary E Goodman^43^, Angela Cox^44^, Fiona Taylor^44^, Penella Woll^44^, Irene Brüske^45^, Heinrich Wichmann^45–47^, Judith Manz^48^, Thomas R Muley^49,50^, Angela Risch^48–52^, Albert Rosenberger^12^, Kjell Grankvist^53^, Mikael Johansson^54^, Frances A shepherd^30^, Ming-Sound Tsao^30^, Susanne M Arnold^55^, Eric B Haura^56^, Ciprian Bolca^57^, Ivana Holcatova^58^, Vladimir Janout^59^, Milica Kontic^60,61^, Jolanta lissowska^62^, Anush mukeria^63^, Simona Ognjanovic^64^, Tadeusz M Orlowski^65^, Ghislaine Scelo^1^, Beata Swiatkowska^66^, David Zaridze^63^, Per Bakke^67^, Vidar Skaug^21^, Shanbeh Zienolddiny^21^, Eric J Duell^68^, Lesley M Butler^27^, Woon-Puay Koh^69^, Yu-tang Gao^70^, Richard S Houlston^71^, John Mclaughlin^72^, Victoria L Stevens^73^, Philippe Joubert^74^, Maxime Lamontagne^74^, David C Nickle^75^, Ma’en Obeidat^76^ , Wim Timens^77^ , Bin Zhu^5^, Lei Song^5^, Linda Kachuri^2^, María Soler Artigas^78,79^, Martin D Tobin^78,79^, Louise V Wain^78,79^, Spirometa Consortium^80^, Thorunn Rafnar^81^, Thorgeir E Thorgeirsson^81^, Gunnar W Reginsson^81^, Kari Stefansson^81^, Dana B Hancock^82^, Laura J Bierut^83^, Margaret R Spitz^84^, Nathan C Gaddis^85^, Sharon M Lutz^86^, Fangyi Gu^5^, Eric O Johnson^87^, Ahsan Kamal^3^, Claudio Pikielny^3^, Dakai Zhu^3^, Sara Lindströem^88^, Xia Jiang^89^, Rachel F Tyndale^90,91^, Georgia Chenevix-Trench^92^, Jonathan Beesley^92^, Yohan Bossé^74,93^, Stephen Chanock^5^, Paul Brennan^1^, Maria teresa landi^5^, Christopher I Amos^3^

^1^International Agency for Research on Cancer, World Health Organization, Lyon, France. ^2^Lunenfeld-Tanenbaum Research Institute, Sinai Health System, University of Toronto, Toronto, Ontario, Canada. ^3^Biomedical Data Science, Geisel School of Medicine at Dartmouth, Hanover, New Hampshire, USA. ^4^Department of Environmental Health, Harvard T.H. Chan School of Public Health and Massachusetts General Hospital/ Harvard Medical School, Boston, Massachusetts, USA. ^5^Division of Cancer Epidemiology and Genetics, National Cancer Institute, US National Institutes of Health, Bethesda, Maryland, USA. ^6^Centre for Cancer Genetic Epidemiology, University of Cambridge, Cambridge, UK. ^7^Department of Clinical Biochemistry, Herlev and Gentofte Hospital, Copenhagen University Hospital, Copenhagen, Denmark. ^8^Faculty of Health and Medical Sciences, University of Copenhagen, Copenhagen, Denmark. ^9^Copenhagen General Population Study, Herlev and Gentofte Hospital, Copenhagen, Denmark. ^10^Department of Epidemiology, University of Texas MD Anderson Cancer Center, Houston, Texas, USA. ^11^Epidemiology Program, University of Hawaii Cancer Center, Honolulu, Hawaii, USA. ^12^Department of Genetic Epidemiology, University Medical Center, Georg August University Göttingen, Göttingen, Germany. ^13^Department of Thoracic Surgery, Division of Epidemiology, Vanderbilt University Medical Center, Nashville, Tennessee, USA. ^14^Department of Epidemiology and Biostatistics, School of Medicine, Case Western Reserve University, Cleveland, Ohio, USA. ^15^University of Oviedo and CIBERESP, Faculty of Medicine, Oviedo, Spain. 16Clalit National Cancer Control Center at Carmel Medical Center and Technion Faculty of Medicine, Haifa, Israel. ^17^School of Health and Related Research, University of Sheffield, Sheffield, UK. ^18^Institute of Translational Medicine, University of Liverpool, Liverpool, UK. ^19^Departments of Health Evidence and Urology, Radboud University Medical Center, Nijmegen, the Netherlands. ^20^Department of Pharmaceutical Sciences, College of Pharmacy, Washington State University, Spokane, Washington, USA. ^21^National Institute of Occupational Health, Oslo, Norway. ^22^British Columbia Cancer Agency, Vancouver, British Columbia, Canada. ^23^Department of Cancer Epidemiology, H. Lee Moffitt Cancer Center and Research Institute, Tampa, Florida, USA. ^24^Department of Epidemiology, Geisel School of Medicine, Hanover, New Hampshire, USA. ^25^Department of Epidemiology and Biostatistics, Jiangsu Key Lab of Cancer Biomarkers, Prevention and Treatment, Collaborative Innovation Center for Cancer Personalized Medicine, School of Public Health, Nanjing Medical University, Nanjing, China. ^26^Department of Preventive Medicine, Seoul National University College of Medicine, Seoul, Republic of Korea. ^27^University of Pittsburgh Cancer Institute, Pittsburgh, Pennsylvania, USA. ^28^Department of Preventive Medicine, IRCCS Foundation Cà Granda Ospedale Maggiore Policlinico, Milan, Italy. ^29^Department of Clinical Sciences and Community Health–DISCCO, University of Milan, Milan, Italy. ^30^University Health Network, Princess Margaret Cancer Centre, Toronto, Ontario, Canada. ^31^Department of Oncology, Herlev and Gentofte Hospital, Copenhagen University Hospital, Copenhagen, Denmark. ^32^Department of Preventive Medicine, Keck School of Medicine, University of Southern California Norris Comprehensive Cancer Center, Los Angeles, California, USA. ^33^Department of Integrative Bioscience and Biotechnology, Sejong University, Seoul, Republic of Korea. ^34^Department of Pathology, Lund University, Lund, Sweden. ^35^Faculty of Medicine, Lund University, Lund, Sweden. ^36^School of Public Health, St Mary’s Campus, Imperial College London, London, UK. ^37^Section for Epidemiology, Department of Public Health, Aarhus University, Aarhus, Denmark. ^38^Hellenic Health Foundation, Athens, Greece. 39Molecular and Nutritional Epidemiology Unit, CSPO (Cancer Research and Prevention Centre), Scientific Institute of Tuscany, Florence, Italy. ^40^Program in Epidemiology, Fred Hutchinson Cancer Research Center, Seattle, Washington, USA. ^41^Huntsman Cancer Institute, Salt Lake City, Utah, USA. ^42^Huntsman Cancer Institute, Department of Population Health Sciences, University of Utah, Salt Lake City, Utah, USA. ^43^Swedish Medical Group, Seattle, Washington, USA. ^44^Department of Oncology, University of Sheffield, Sheffield, UK. ^45^Institute of Epidemiology II, Helmholtz Zentrum München–German Research Center for Environmental Health, Neuherberg, Germany. ^46^Institute of Medical Informatics, Biometry and Epidemiology, Ludwig Maximilians University, Munich, Germany. ^47^Institute of Medical Statistics and Epidemiology, Technical University of Munich, Munich, Germany. ^48^Research Unit of Molecular Epidemiology, Helmholtz Zentrum München–German Research Center for Environmental Health, Neuherberg, Germany. ^49^Thoraxklinik at University Hospital Heidelberg, Heidelberg, Germany. ^50^Translational Lung Research Center Heidelberg (TLRC-H), Heidelberg, Germany. ^51^German Center for Lung Research (DZL), Heidelberg, Germany. ^52^University of Salzburg and Cancer Cluster Salzburg, Salzburg, Austria. ^53^Department of Medical Biosciences, Umeå University, Umeå, Sweden. ^54^Department of Radiation Sciences, Umeå University, Umeå, Sweden. ^55^Markey Cancer Center, University of Kentucky, Lexington, Kentucky, USA. ^56^Department of Thoracic Oncology, H. Lee Moffitt Cancer Center and Research Institute, Tampa, Florida, USA. ^57^Institute of Pneumology ‘Marius Nasta’, Bucharest, Romania. ^58^2nd Faculty of Medicine, Charles University, Prague, Czech Republic. ^59^Faculty of Medicine, University of Ostrava, Ostrava, Czech Republic. ^60^Clinical Center of Serbia, Belgrade, Serbia. ^61^School of Medicine, University of Belgrade, Belgrade, Serbia. ^62^M. Sklodowska-Curie Cancer Center, Institute of Oncology, Warsaw, Poland. ^63^Department of Epidemiology and Prevention, Russian N.N.Blokhin Cancer Research Centre, Moscow, Russian Federation. ^64^International Organization for Cancer Prevention and Research, Belgrade, Serbia. ^65^Department of Surgery, National Tuberculosis and Lung Diseases Research Institute, Warsaw, Poland. ^66^Nofer Institute of Occupational Medicine, Department of Environmental Epidemiology, Lodz, Poland. ^67^Department of Clinical Science, University of Bergen, Bergen, Norway. ^68^Unit of Nutrition and Cancer, Catalan Institute of Oncology (ICO-IDIBELL), Barcelona, Spain. ^69^Duke–National University of Singapore Medical School, Singapore. ^70^Department of Epidemiology, Shanghai Cancer Institute, Shanghai, China. ^71^The Institute of Cancer Research, London, UK. ^72^Public Health Ontario, Toronto, Ontario, Canada. ^73^American Cancer Society, Atlanta, Georgia, USA. ^74^Institut Universitaire de Cardiologie et de Pneumologie de Québec, Québec, Québec, Canada. ^75^Merck Research Laboratories, Genetics and Pharmacogenomics, Boston, Massachusetts, USA. ^76^University of British Columbia Centre for Heart Lung Innovation, St Paul’s Hospital, Vancouver, British Columbia, Canada. ^77^University of Groningen, University Medical Center Groningen, Department of Pathology and Medical Biology, GRIAC Research Institute, Groningen, the Netherlands. ^78^Genetic Epidemiology Group, Department of Health Sciences, University of Leicester, Leicester, UK. ^79^National Institute for Health Research (NIHR) Leicester Respiratory Biomedical Research Unit, Glenfield Hospital, Leicester, UK. ^80^A list of members appears in the supplementary Note. ^81^deCODE Genetics, Amgen, Inc., Reykjavik, Iceland. ^82^Behavioral and Urban Health Program, Behavioral Health and Criminal Justice Division, RTI International, Research Triangle Park, North Carolina, USA. ^83^Department of Psychiatry, Washington University School of Medicine, St. Louis, Missouri, USA. ^84^Duncan Cancer Center, Baylor College of Medicine, Houston, Texas, USA. ^85^Research Computing Division, RTI International, Research Triangle Park, North Carolina, USA. ^86^Department of Biostatistics and Informatics, University of Colorado Anschutz Medical Campus, Aurora, Colorado, USA. ^87^Fellow Program and Behavioral Health and Criminal Justice Division, RTI International, Research Triangle Park, North Carolina, USA. ^88^Department of Epidemiology, University of Washington, Seattle, Washington, USA. ^89^Department of Epidemiology, Harvard T.H. Chan School of Public Health, Boston, Massachusetts, USA. ^90^Departments of Pharmacology and Toxicology & Psychiatry, University of Toronto, Toronto, Ontario, Canada. ^91^Campbell Family Mental Health Research Institute, Centre for Addiction and Mental Health, Toronto, Ontario, Canada. ^92^Cancer Division, QIMR Berghofer Medical Research Institute, Brisbane, Queensland, Australia. ^93^Department of Molecular Medicine, Laval University, Québec, Québec, Canada.

Supplementary Table 1	Association of genome-wide significant SNPs for self-reported moderate-to-vigorous physical activity from the GWAS by Klimentidis et al. (1) with overall lung cancer

| SNP | CHR | Position (hg19/b37) | EA | OA | BETA | SE | *P*-value |
| --- | --- | --- | --- | --- | --- | --- | --- |
| rs2942127 | 1 | 204420067 | G | A | -0.035 | 0.015 | 0.018 |
| rs1974771 | 2 | 54278543 | G | A | -0.014 | 0.019 | 0.441 |
| rs2114286 | 3 | 41194283 | A | G | -0.013 | 0.018 | 0.474 |
| rs877483 | 3 | 53846741 | T | C | 0.013 | 0.015 | 0.383 |
| rs2035562 | 3 | 85056521 | A | G | 0.015 | 0.012 | 0.232 |
| rs1972763 | 4 | 159860563 | C | T | 0.000 | 0.012 | 0.987 |
| rs77742115 | 5 | 18330424 | T | C | -0.008 | 0.017 | 0.650 |
| rs1186721 | 7 | 34974602 | G | A | 0.014 | 0.013 | 0.276 |
| rs921915 | 7 | 50228581 | T | C | 0.007 | 0.012 | 0.561 |
| rs1043595 | 7 | 128410012 | G | A | -0.005 | 0.020 | 0.796 |
| rs7804463 | 7 | 133447651 | T | C | 0.002 | 0.014 | 0.861 |
| rs2988004 | 9 | 37044388 | T | G | 0.016 | 0.012 | 0.190 |
| rs7326482 | 13 | 54037803 | G | T | -0.007 | 0.012 | 0.552 |
| rs10145335 | 14 | 98547748 | G | A | 0.000 | 0.014 | 0.995 |
| rs4886868 | 15 | 74353561 | T | G | -0.009 | 0.013 | 0.485 |
| rs12912808 | 15 | 95292223 | C | T | -0.017 | 0.019 | 0.385 |
| rs429358 | 19 | 45411941 | T | C | 0.041 | 0.018 | 0.019 |
| rs1921981 | 21 | 42422547 | G | A | -0.015 | 0.017 | 0.367 |

EA, effect allele. OA, other allele. EAF, effect allele frequency. SE, standard error.

Supplementary Table 2	Association of genome-wide significant SNPs for accelerometer-measured physical activity with overall lung cancer

| SNP | CHR | Position (hg19/b37) | EA | OA | BETA | SE | *P*-value |
| --- | --- | --- | --- | --- | --- | --- | --- |
| Primary analyis (SNPs from GWAS by Klimentidits et al. (1) – ‘average acceleration’ | | | | | | | |
| rs336605 | 3 | 18656350 | G | T | -0.033 | 0.021 | 0.113 |
| rs10067451 | 5 | 87942506 | G | A | -0.001 | 0.018 | 0.945 |
| rs28749810 | 5 | 152048630 | C | A | 0.009 | 0.014 | 0.507 |
| rs7084454 | 10 | 21821274 | G | A | -0.003 | 0.012 | 0.839 |
| rs148193266 | 11 | 104528681 | A | C | -0.020 | 0.039 | 0.615 |
| rs79724577 | 17 | 43463493 | A | C | 0.022 | 0.015 | 0.152 |
| rs1518139 | 18 | 40751232 | G | T | 0.005 | 0.015 | 0.715 |
| Secondary analyis (SNPs from GWAS by Doherty et al. (2) – ‘overall activity’ | | | | | | | |
| rs6775319 | 3 | 18758501 | A | T | -0.030 | 0.021 | 0.152 |
| rs9293503 | 5 | 87948962 | T | C | 0.000 | 0.018 | 0.986 |
| rs6873698 | 5 | 152039420 | C | T | 0.012 | 0.014 | 0.387 |
| rs11012732 | 10 | 21830104 | A | G | -0.005 | 0.012 | 0.674 |
| rs59499656 | 18 | 40768309 | A | T | 0.007 | 0.015 | 0.653 |

EA, effect allele. OA, other allele. EAF, effect allele frequency. SE, standard error.

Supplementary Table 3	Association (P<5x10^-8^) of the 19 SNPs used as candidate genetic instruments for moderate-to-vigorous physical activity with confounders or lung cancer, in the GWAS by Klimentidis et al. (1)

| SNP | CHR | Position (hg19/b37) | Trait | Excluded from MR analysis |
| --- | --- | --- | --- | --- |
| rs2942127 | 1 | 204420067 | None | No |
| rs1974771 | 2 | 54278543 | None | No |
| rs2114286 | 3 | 41194283 | None | No |
| rs877483 | 3 | 53846741 | None | No |
| rs2035562 | 3 | 85056521 | None | No |
| rs1972763 | 4 | 159860563 | None | No |
| rs77742115 | 5 | 18330424 | None | No |
| rs2854277 | 6 | 32628084 | Lung cancer (PMID: 22899653) Squamous cell lung carcinoma (PMID: 28604730), Lung cancer in ever smokers (PMID: 28604730) | Yes |
| rs1186721 | 7 | 34974602 | None | No |
| rs921915 | 7 | 50228581 | None | No |
| rs1043595 | 7 | 128410012 | None | No |
| rs7804463 | 7 | 133447651 | None | No |
| rs2988004 | 9 | 37044388 | None | No |
| rs7326482 | 13 | 54037803 | None | No |
| rs10145335 | 14 | 98547748 | None | No |
| rs4886868 | 15 | 74353561 | None | No |
| rs12912808 | 15 | 95292223 | None | No |
| rs429358 | 19 | 45411941 | None | No |
| rs1921981 | 21 | 42422547 | None | No |

Supplementary Table 4	Association (P<5x10^-8^) of the 19 SNPs used as candidate genetic instruments for accelerometer-measured physical activity with confounders or lung cancer

| SNP | CHR | Position (hg19/b37) | Trait | Excluded from MR analysis |
| --- | --- | --- | --- | --- |
| Primary analyis (SNPs from GWAS by Klimentidits et al. (1)) | | | | |
| rs34517439 | 1 | 78450517 | Lung cancer (PMID: 28604730), Forced expiratory volume in 1-second (Neale Lab - UK Biobank) | Yes |
| rs336605 | 3 | 18656350 | None | No |
| rs10067451 | 5 | 87942506 | None | No |
| rs28749810 | 5 | 152048630 | None | No |
| rs7084454 | 10 | 21821274 | None | No |
| rs148193266 | 11 | 104528681 | None | No |
| rs79724577 | 17 | 43463493 | None | No |
| rs1518139 | 18 | 40751232 | None | No |
| Secondary analyis (SNPs from GWAS by Doherty et al. (2)) | | | | |
| rs6775319 | 3 | 18758501 | None | No |
| rs9293503 | 5 | 87948962 | None | No |
| rs6873698 | 5 | 152039420 | None | No |
| rs11012732 | 10 | 21830104 | None | No |
| rs2696625 | 17 | 44326864 | Forced vital capacity (Neale Lab - UK Biobank) | Yes |
| rs59499656 | 18 | 40768309 | None | No |

Supplementary Table 5	Sampe size and a priori power estimates

| Cancer type | Exposure | Sample size for outcome | Proportion of cases | R² | OR=0.90 | OR=0.85 | OR=0.80 | OR=0.75 | OR=0.70 | OR=0.65 |
| --- | --- | --- | --- | --- | --- | --- | --- | --- | --- | --- |
| Overall lung cancer | Self-reported moderate-to-vigorous | 85716 | 0.54 | 0.007 | 0.26 | 0.52 | 0.80 | 0.95 | 0.99 | 1.00 |
|  | Accelerometer-measured (‘average acceleration’) | 85716 | 0.54 | 0.003 | 0.12 | 0.23 | 0.39 | 0.58 | 0.77 | 0.91 |
|  | Accelerometer-measured (‘overall activity’) | 85716 | 0.54 | 0.002 | 0.10 | 0.18 | 0.30 | 0.46 | 0.63 | 0.80 |
| Adenocarcinoma | Self-reported moderate-to-vigorous physical activity | 66756 | 0.32 | 0.009 | 0.22 | 0.45 | 0.70 | 0.89 | 0.97 | 1.00 |
|  | Accelerometer-measured (‘average acceleration’) | 66756 | 0.32 | 0.003 | 0.11 | 0.20 | 0.32 | 0.48 | 0.65 | 0.80 |
|  | Accelerometer-measured (‘overall activity’) | 66756 | 0.32 | 0.002 | 0.09 | 0.16 | 0.25 | 0.37 | 0.52 | 0.67 |
| Squamous cell carcinoma | Self-reported moderate-to-vigorous physical activity | 63053 | 0.23 | 0.010 | 0.19 | 0.37 | 0.59 | 0.79 | 0.92 | 0.98 |
|  | Accelerometer-measured (‘average acceleration’) | 63053 | 0.23 | 0.004 | 0.10 | 0.16 | 0.26 | 0.39 | 0.53 | 0.68 |
|  | Accelerometer-measured (‘overall activity’) | 63053 | 0.23 | 0.003 | 0.09 | 0.13 | 0.20 | 0.30 | 0.41 | 0.54 |
| Small cell carcinoma | Self-reported moderate-to-vigorous physical activity | 24108 | 0.11 | 0.025 | 0.13 | 0.23 | 0.37 | 0.53 | 0.69 | 0.83 |
|  | Accelerometer-measured (‘average acceleration’) | 24108 | 0.11 | 0.009 | 0.08 | 0.11 | 0.16 | 0.23 | 0.32 | 0.42 |
|  | Accelerometer-measured (‘overall activity’) | 24108 | 0.11 | 0.007 | 0.07 | 0.09 | 0.13 | 0.18 | 0.24 | 0.32 |
| Never smoker | Self-reported moderate-to-vigorous physical activity | 9859 | 0.30 | 0.062 | 0.21 | 0.43 | 0.68 | 0.87 | 0.96 | 0.99 |
|  | Accelerometer-measured (‘average acceleration’) | 9859 | 0.30 | 0.023 | 0.11 | 0.19 | 0.31 | 0.46 | 0.62 | 0.77 |
|  | Accelerometer-measured (‘overall activity’) | 9859 | 0.30 | 0.016 | 0.09 | 0.15 | 0.24 | 0.35 | 0.49 | 0.63 |
| Ever smoker | Self-reported moderate-to-vigorous physical activity | 40187 | 0.62 | 0.015 | 0.25 | 0.51 | 0.79 | 0.95 | 0.99 | 1.00 |
|  | Accelerometer-measured (‘average acceleration’) | 40187 | 0.62 | 0.006 | 0.12 | 0.22 | 0.38 | 0.58 | 0.77 | 0.91 |
|  | Accelerometer-measured (‘overall activity’) | 40187 | 0.62 | 0.004 | 0.10 | 0.18 | 0.29 | 0.45 | 0.64 | 0.81 |

R², explained variation by SNPs. OR, odds ratio. ^a^ Primary analysis used GWAS data from Klementidits et al. (1). ^b^Secondary analysis used GWAS data from Doherty et al. (2).

Supplementary Table 6	Mendelian randomization estimates for the relationship between accelerometer-measured physical activity (‘overall activty’) and lung cancer

| Outcomes | Method | OR^a^ | (95% CI)^a^ | *P*-value | *Q*-Value |
| --- | --- | --- | --- | --- | --- |
| Overall lung cancer | Inverse-variance weighted | 0.89 | (0.56;1.41) | 0.619 | 0.816 |
|  | Maximum likelihood | 0.89 | (0.56;1.41) | 0.618 | 0.760 |
|  | Weighted median | 0.87 | (0.49;1.55) | 0.643 | 0.813 |
|  | MR PRESSO | 0.89 | (0.6;1.32) | 0.593 | 0.813 |
| Adenocarcinoma | Inverse-variance weighted | 0.66 | (0.37;1.19) | 0.164 | 0.429 |
|  | Maximum likelihood | 0.65 | (0.36;1.19) | 0.163 | 0.645 |
|  | Weighted median | 0.67 | (0.32;1.41) | 0.295 | 0.427 |
|  | MR PRESSO | 0.66 | (0.38;1.14) | 0.211 | 0.427 |
| Squamous cell carcinoma | Inverse-variance weighted | 1.06 | (0.49;2.29) | 0.874 | 0.874 |
|  | Maximum likelihood | 1.07 | (0.49;2.31) | 0.873 | 0.760 |
|  | Weighted median | 1.17 | (0.44;3.12) | 0.760 | 0.873 |
|  | MR PRESSO | 1.06 | (0.5;2.26) | 0.880 | 0.873 |
| Small cell carcinoma | Inverse-variance weighted | 2.33 | (0.61;8.85) | 0.214 | 0.429 |
|  | Maximum likelihood | 2.35 | (0.61;9.07) | 0.214 | 0.645 |
|  | Weighted median | 2.28 | (0.45;11.68) | 0.323 | 0.427 |
|  | MR PRESSO | 2.33 | (0.86;6.33) | 0.173 | 0.427 |
| Never smoker | Inverse-variance weighted | 0.40 | (0.11;1.42) | 0.154 | 0.429 |
|  | Maximum likelihood | 0.39 | (0.11;1.42) | 0.153 | 0.645 |
|  | Weighted median | 0.29 | (0.05;1.51) | 0.140 | 0.427 |
|  | MR PRESSO | 0.40 | (0.13;1.22) | 0.181 | 0.427 |
| Ever smoker | Inverse-variance weighted | 0.89 | (0.52;1.54) | 0.680 | 0.816 |
|  | Maximum likelihood | 0.89 | (0.51;1.54) | 0.677 | 0.760 |
|  | Weighted median | 0.83 | (0.42;1.65) | 0.601 | 0.813 |
|  | MR PRESSO | 0.89 | (0.53;1.51) | 0.690 | 0.813 |

MR PRESSO, MR Pleiotropy RESidual Sum and Outlier. ^a^ OR (odds ratio) per one standard deviation increment in ‘overall activity’.

Supplementary Table 7	Between SNP-heterogeneity

| Outcome | Exposure | Cochran’s Q | Degrees of Freedom | *P*-value |
| --- | --- | --- | --- | --- |
| Overall lung cancer | Self-reported moderate-to-vigorous | 17.3 | 17 | 0.437 |
|  | Accelerometer-measured (‘average acceleration’) | 4.6 | 6 | 0.590 |
|  | Accelerometer-measured (‘overall activity’) | 2.9 | 4 | 0.570 |
| Adenocarcinoma | Self-reported moderate-to-vigorous | 19.8 | 16 | 0.230 |
|  | Accelerometer-measured (‘average acceleration’) | 5.8 | 6 | 0.449 |
|  | Accelerometer-measured (‘overall activity’) | 3.5 | 4 | 0.480 |
| Squamous cell carcinoma | Self-reported moderate-to-vigorous | 23.1 | 17 | 0.146 |
|  | Accelerometer-measured (‘average acceleration’) | 5.1 | 6 | 0.533 |
|  | Accelerometer-measured (‘overall activity’) | 3.9 | 4 | 0.424 |
| Small cell carcinoma | Self-reported moderate-to-vigorous | 19.7 | 17 | 0.289 |
|  | Accelerometer-measured (‘average acceleration’) | 5.1 | 5 | 0.406 |
|  | Accelerometer-measured (‘overall activity’) | 2.2 | 4 | 0.691 |
| Never smoker | Self-reported moderate-to-vigorous | 19.4 | 17 | 0.305 |
|  | Accelerometer-measured (‘average acceleration’) | 3.2 | 6 | 0.786 |
|  | Accelerometer-measured (‘overall activity’) | 3.1 | 4 | 0.540 |
| Ever smoker | Self-reported moderate-to-vigorous | 9.7 | 17 | 0.916 |
|  | Accelerometer-measured (‘average acceleration’) | 4.5 | 6 | 0.615 |
|  | Accelerometer-measured (‘overall activity’) | 3.7 | 4 | 0.447 |

Supplementary Table 8	Inverse variance weighted estimates for self-reported moderate-to-vigorous physical activity and lung cancer, with SNPs individually removed in leave-one-out analyses

| Outcome | SNP excluded | OR^a^ | (95% CI)^a^ | *P*-value | *Q*-value |
| --- | --- | --- | --- | --- | --- |
| Overall lung cancer | rs10145335 | 0.65 | (0.4;1.05) | 0.079 | 0.160 |
|  | rs1043595 | 0.66 | (0.41;1.07) | 0.094 | 0.163 |
|  | rs1186721 | 0.69 | (0.43;1.12) | 0.132 | 0.163 |
|  | rs12912808 | 0.68 | (0.42;1.1) | 0.119 | 0.163 |
|  | rs1921981 | 0.68 | (0.42;1.1) | 0.115 | 0.163 |
|  | rs1972763 | 0.65 | (0.4;1.05) | 0.080 | 0.160 |
|  | rs1974771 | 0.61 | (0.38;0.98) | 0.042 | 0.160 |
|  | rs2035562 | 0.70 | (0.43;1.13) | 0.142 | 0.163 |
|  | rs2114286 | 0.64 | (0.4;1.02) | 0.060 | 0.160 |
|  | rs2942127 | 0.75 | (0.47;1.2) | 0.226 | 0.239 |
|  | rs2988004 | 0.70 | (0.44;1.13) | 0.145 | 0.163 |
|  | rs429358 | 0.76 | (0.47;1.22) | 0.260 | 0.260 |
|  | rs4886868 | 0.62 | (0.39;1) | 0.049 | 0.160 |
|  | rs7326482 | 0.62 | (0.39;1) | 0.050 | 0.160 |
|  | rs77742115 | 0.63 | (0.39;1.01) | 0.056 | 0.160 |
|  | rs7804463 | 0.64 | (0.4;1.04) | 0.070 | 0.160 |
|  | rs877483 | 0.63 | (0.4;1) | 0.049 | 0.160 |
|  | rs921915 | 0.67 | (0.41;1.09) | 0.109 | 0.163 |
| Adenocarcinoma | rs10145335 | 0.72 | (0.34;1.5) | 0.375 | 0.732 |
|  | rs1043595 | 0.82 | (0.39;1.72) | 0.593 | 0.732 |
|  | rs1186721 | 0.82 | (0.39;1.71) | 0.590 | 0.732 |
|  | rs12912808 | 0.87 | (0.43;1.77) | 0.702 | 0.732 |
|  | rs1921981 | 0.89 | (0.44;1.77) | 0.732 | 0.732 |
|  | rs1972763 | 0.80 | (0.38;1.68) | 0.557 | 0.732 |
|  | rs1974771 | 0.66 | (0.33;1.31) | 0.234 | 0.732 |
|  | rs2035562 | 0.79 | (0.37;1.68) | 0.546 | 0.732 |
|  | rs2114286 | 0.75 | (0.36;1.59) | 0.458 | 0.732 |
|  | rs2942127 | 0.76 | (0.36;1.6) | 0.467 | 0.732 |
|  | rs2988004 | 0.75 | (0.36;1.56) | 0.443 | 0.732 |
|  | rs429358 | 0.86 | (0.41;1.8) | 0.687 | 0.732 |
|  | rs4886868 | 0.74 | (0.35;1.55) | 0.424 | 0.732 |
|  | rs7326482 | 0.77 | (0.36;1.63) | 0.492 | 0.732 |
|  | rs77742115 | 0.63 | (0.33;1.21) | 0.167 | 0.732 |
|  | rs877483 | 0.71 | (0.35;1.43) | 0.335 | 0.732 |
|  | rs921915 | 0.84 | (0.4;1.76) | 0.645 | 0.732 |
| Squamous cell carcinoma | rs10145335 | 0.46 | (0.19;1.12) | 0.087 | 0.125 |
|  | rs1043595 | 0.43 | (0.18;1.02) | 0.057 | 0.125 |
|  | rs1186721 | 0.51 | (0.22;1.2) | 0.121 | 0.145 |
|  | rs12912808 | 0.44 | (0.18;1.06) | 0.067 | 0.125 |
|  | rs1921981 | 0.48 | (0.2;1.16) | 0.104 | 0.134 |
|  | rs1972763 | 0.43 | (0.18;1.05) | 0.065 | 0.125 |
|  | rs1974771 | 0.40 | (0.17;0.95) | 0.038 | 0.125 |
|  | rs2035562 | 0.46 | (0.19;1.1) | 0.081 | 0.125 |
|  | rs2114286 | 0.36 | (0.17;0.77) | 0.009 | 0.125 |
|  | rs2942127 | 0.52 | (0.22;1.21) | 0.130 | 0.145 |
|  | rs2988004 | 0.53 | (0.23;1.22) | 0.137 | 0.145 |
|  | rs429358 | 0.57 | (0.25;1.28) | 0.173 | 0.173 |
|  | rs4886868 | 0.45 | (0.19;1.07) | 0.072 | 0.125 |
|  | rs7326482 | 0.39 | (0.17;0.91) | 0.028 | 0.125 |
|  | rs77742115 | 0.45 | (0.18;1.09) | 0.076 | 0.125 |
|  | rs7804463 | 0.40 | (0.16;0.95) | 0.039 | 0.125 |
|  | rs877483 | 0.46 | (0.19;1.13) | 0.090 | 0.125 |
|  | rs921915 | 0.46 | (0.19;1.12) | 0.086 | 0.125 |
| Small cell carcinoma | rs10145335 | 0.32 | (0.08;1.29) | 0.109 | 0.245 |
|  | rs1043595 | 0.35 | (0.09;1.45) | 0.149 | 0.245 |
|  | rs1186721 | 0.37 | (0.09;1.56) | 0.178 | 0.245 |
|  | rs12912808 | 0.39 | (0.09;1.62) | 0.194 | 0.245 |
|  | rs1921981 | 0.53 | (0.14;1.92) | 0.332 | 0.352 |
|  | rs1972763 | 0.37 | (0.09;1.57) | 0.179 | 0.245 |
|  | rs1974771 | 0.36 | (0.09;1.45) | 0.149 | 0.245 |
|  | rs2035562 | 0.41 | (0.1;1.62) | 0.204 | 0.245 |
|  | rs2114286 | 0.34 | (0.09;1.35) | 0.126 | 0.245 |
|  | rs2942127 | 0.45 | (0.12;1.67) | 0.232 | 0.261 |
|  | rs2988004 | 0.37 | (0.09;1.57) | 0.179 | 0.245 |
|  | rs429358 | 0.61 | (0.16;2.26) | 0.456 | 0.456 |
|  | rs4886868 | 0.28 | (0.08;1.05) | 0.059 | 0.245 |
|  | rs7326482 | 0.34 | (0.09;1.36) | 0.127 | 0.245 |
|  | rs77742115 | 0.36 | (0.09;1.54) | 0.169 | 0.245 |
|  | rs7804463 | 0.31 | (0.07;1.3) | 0.109 | 0.245 |
|  | rs877483 | 0.36 | (0.09;1.47) | 0.154 | 0.245 |
|  | rs921915 | 0.33 | (0.08;1.28) | 0.109 | 0.245 |
| Never smoker | rs10145335 | 0.78 | (0.18;3.48) | 0.747 | 0.747 |
|  | rs1043595 | 0.67 | (0.14;3.26) | 0.620 | 0.695 |
|  | rs1186721 | 0.57 | (0.11;2.9) | 0.494 | 0.656 |
|  | rs12912808 | 0.48 | (0.09;2.45) | 0.376 | 0.656 |
|  | rs1921981 | 0.55 | (0.11;2.71) | 0.459 | 0.656 |
|  | rs1972763 | 0.34 | (0.08;1.51) | 0.157 | 0.656 |
|  | rs1974771 | 0.60 | (0.12;3.06) | 0.539 | 0.656 |
|  | rs2035562 | 0.70 | (0.15;3.33) | 0.656 | 0.695 |
|  | rs2114286 | 0.47 | (0.1;2.31) | 0.354 | 0.656 |
|  | rs2942127 | 0.51 | (0.1;2.66) | 0.424 | 0.656 |
|  | rs2988004 | 0.55 | (0.11;2.73) | 0.468 | 0.656 |
|  | rs429358 | 0.50 | (0.1;2.46) | 0.391 | 0.656 |
|  | rs4886868 | 0.44 | (0.09;2.17) | 0.312 | 0.656 |
|  | rs7326482 | 0.41 | (0.08;1.96) | 0.261 | 0.656 |
|  | rs77742115 | 0.51 | (0.1;2.56) | 0.417 | 0.656 |
|  | rs7804463 | 0.34 | (0.07;1.6) | 0.172 | 0.656 |
|  | rs877483 | 0.51 | (0.1;2.53) | 0.408 | 0.656 |
|  | rs921915 | 0.60 | (0.12;3.11) | 0.547 | 0.656 |
| Ever smoker | rs10145335 | 0.64 | (0.34;1.21) | 0.171 | 0.477 |
|  | rs1043595 | 0.71 | (0.38;1.33) | 0.289 | 0.477 |
|  | rs1186721 | 0.74 | (0.39;1.41) | 0.363 | 0.477 |
|  | rs12912808 | 0.75 | (0.4;1.41) | 0.371 | 0.477 |
|  | rs1921981 | 0.79 | (0.42;1.48) | 0.457 | 0.514 |
|  | rs1972763 | 0.71 | (0.37;1.34) | 0.290 | 0.477 |
|  | rs1974771 | 0.69 | (0.36;1.31) | 0.253 | 0.477 |
|  | rs2035562 | 0.73 | (0.39;1.38) | 0.334 | 0.477 |
|  | rs2114286 | 0.70 | (0.37;1.32) | 0.274 | 0.477 |
|  | rs2942127 | 0.82 | (0.44;1.55) | 0.543 | 0.543 |
|  | rs2988004 | 0.78 | (0.41;1.48) | 0.445 | 0.514 |
|  | rs429358 | 0.81 | (0.42;1.55) | 0.526 | 0.543 |
|  | rs4886868 | 0.71 | (0.38;1.35) | 0.299 | 0.477 |
|  | rs7326482 | 0.73 | (0.39;1.37) | 0.334 | 0.477 |
|  | rs77742115 | 0.71 | (0.37;1.34) | 0.286 | 0.477 |
|  | rs7804463 | 0.74 | (0.39;1.38) | 0.341 | 0.477 |
|  | rs877483 | 0.70 | (0.37;1.32) | 0.268 | 0.477 |
|  | rs921915 | 0.72 | (0.38;1.37) | 0.310 | 0.477 |

OR, odds ratio. CI, confidence interval.

Supplementary Table 9	Inverse variance weighted estimates for accelerometer-based (‘average acceleration’) physical activity and lung cancer, with SNPs individually removed in leave-one-out analyses

| Outcome | SNP excluded | OR^a^ | (95% CI)^a^ | *P*-value | *Q*-value |
| --- | --- | --- | --- | --- | --- |
| Overall lung cancer | rs11012732 | 0.96 | (0.91;1.03) | 0.248 | 0.363 |
|  | rs12522261 | 0.95 | (0.91;1.01) | 0.081 | 0.363 |
|  | rs148193266 | 0.96 | (0.91;1.01) | 0.135 | 0.363 |
|  | rs56194509 | 0.99 | (0.94;1.04) | 0.741 | 0.741 |
|  | rs59499656 | 0.97 | (0.91;1.03) | 0.259 | 0.363 |
|  | rs6775319 | 0.97 | (0.92;1.03) | 0.318 | 0.371 |
|  | rs9293503 | 0.96 | (0.9;1.02) | 0.178 | 0.363 |
| Adenocarcinoma | rs11012732 | 0.95 | (0.88;1.04) | 0.269 | 0.335 |
|  | rs12522261 | 0.93 | (0.86;1.01) | 0.070 | 0.191 |
|  | rs148193266 | 0.93 | (0.87;1) | 0.054 | 0.191 |
|  | rs56194509 | 0.97 | (0.9;1.03) | 0.319 | 0.335 |
|  | rs59499656 | 0.93 | (0.86;1.01) | 0.082 | 0.191 |
|  | rs6775319 | 0.96 | (0.89;1.03) | 0.242 | 0.335 |
|  | rs9293503 | 0.96 | (0.89;1.04) | 0.335 | 0.335 |
| Squamous cell carcinoma | rs11012732 | 1.03 | (0.94;1.12) | 0.591 | 0.948 |
|  | rs12522261 | 1.02 | (0.93;1.11) | 0.666 | 0.948 |
|  | rs148193266 | 1.01 | (0.92;1.1) | 0.891 | 0.948 |
|  | rs56194509 | 1.02 | (0.93;1.11) | 0.713 | 0.948 |
|  | rs59499656 | 1.03 | (0.94;1.12) | 0.557 | 0.948 |
|  | rs6775319 | 1.05 | (0.96;1.14) | 0.295 | 0.948 |
|  | rs9293503 | 1.00 | (0.92;1.1) | 0.948 | 0.948 |
| Small cell carcinoma | rs11012732 | 1.05 | (0.89;1.23) | 0.572 | 0.917 |
|  | rs12522261 | 1.06 | (0.91;1.24) | 0.462 | 0.917 |
|  | rs148193266 | 1.11 | (0.94;1.3) | 0.212 | 0.917 |
|  | rs59499656 | 1.04 | (0.87;1.26) | 0.647 | 0.917 |
|  | rs6775319 | 0.99 | (0.84;1.17) | 0.917 | 0.917 |
|  | rs9293503 | 1.02 | (0.87;1.19) | 0.811 | 0.917 |
| Never smoker | rs11012732 | 0.87 | (0.74;1.01) | 0.060 | 0.294 |
|  | rs12522261 | 0.91 | (0.78;1.06) | 0.210 | 0.294 |
|  | rs148193266 | 0.90 | (0.78;1.04) | 0.152 | 0.294 |
|  | rs56194509 | 0.91 | (0.78;1.05) | 0.176 | 0.294 |
|  | rs59499656 | 0.94 | (0.81;1.09) | 0.380 | 0.380 |
|  | rs6775319 | 0.90 | (0.78;1.04) | 0.154 | 0.294 |
|  | rs9293503 | 0.92 | (0.79;1.08) | 0.312 | 0.364 |
| Ever smoker | rs11012732 | 0.98 | (0.92;1.04) | 0.557 | 0.653 |
|  | rs12522261 | 0.97 | (0.91;1.03) | 0.258 | 0.653 |
|  | rs148193266 | 0.97 | (0.91;1.03) | 0.318 | 0.653 |
|  | rs56194509 | 0.99 | (0.93;1.05) | 0.777 | 0.777 |
|  | rs59499656 | 0.96 | (0.9;1.02) | 0.169 | 0.653 |
|  | rs6775319 | 0.98 | (0.93;1.04) | 0.560 | 0.653 |
|  | rs9293503 | 0.98 | (0.92;1.04) | 0.527 | 0.653 |

OR, odds ratio. CI, confidence interval.

Supplementary Table 10	Inverse variance weighted estimates for accelerometer-based (‘overall acceleration’) physical activity and lung cancer subtype, with SNPs individually removed in leave-one-out analyses

| Outcome | SNP excluded | OR^a^ | (95% CI)^a^ | *P*-value | *Q*-value |
| --- | --- | --- | --- | --- | --- |
| Overall lung cancer | rs11012732 | 0.91 | (0.53;1.57) | 0.746 | 0.932 |
|  | rs59499656 | 0.92 | (0.55;1.53) | 0.739 | 0.932 |
|  | rs6775319 | 0.99 | (0.61;1.6) | 0.955 | 0.955 |
|  | rs6873698 | 0.78 | (0.47;1.29) | 0.335 | 0.932 |
|  | rs9293503 | 0.86 | (0.51;1.46) | 0.573 | 0.932 |
| Adenocarcinoma | rs11012732 | 0.71 | (0.35;1.43) | 0.339 | 0.424 |
|  | rs59499656 | 0.55 | (0.28;1.08) | 0.084 | 0.213 |
|  | rs6775319 | 0.73 | (0.39;1.34) | 0.305 | 0.424 |
|  | rs6873698 | 0.56 | (0.29;1.08) | 0.085 | 0.213 |
|  | rs9293503 | 0.77 | (0.4;1.5) | 0.447 | 0.447 |
| Squamous cell carcinoma | rs11012732 | 1.08 | (0.38;3.06) | 0.885 | 0.885 |
|  | rs59499656 | 1.11 | (0.44;2.82) | 0.823 | 0.885 |
|  | rs6775319 | 1.40 | (0.61;3.2) | 0.431 | 0.885 |
|  | rs6873698 | 0.93 | (0.37;2.35) | 0.882 | 0.885 |
|  | rs9293503 | 0.85 | (0.36;2.01) | 0.717 | 0.885 |
| Small cell carcinoma | rs11012732 | 2.57 | (0.62;10.64) | 0.191 | 0.318 |
|  | rs59499656 | 3.11 | (0.57;16.84) | 0.188 | 0.318 |
|  | rs6775319 | 1.53 | (0.33;7.17) | 0.590 | 0.590 |
|  | rs6873698 | 2.88 | (0.7;11.78) | 0.141 | 0.318 |
|  | rs9293503 | 1.96 | (0.46;8.38) | 0.364 | 0.455 |
| Never smoker | rs11012732 | 0.23 | (0.05;0.97) | 0.046 | 0.230 |
|  | rs59499656 | 0.56 | (0.14;2.32) | 0.423 | 0.423 |
|  | rs6775319 | 0.37 | (0.09;1.44) | 0.151 | 0.378 |
|  | rs6873698 | 0.43 | (0.1;1.81) | 0.248 | 0.411 |
|  | rs9293503 | 0.48 | (0.11;2.07) | 0.329 | 0.411 |
| Ever smoker | rs11012732 | 0.99 | (0.52;1.91) | 0.984 | 0.984 |
|  | rs59499656 | 0.72 | (0.38;1.36) | 0.311 | 0.984 |
|  | rs6775319 | 0.98 | (0.56;1.73) | 0.948 | 0.984 |
|  | rs6873698 | 0.81 | (0.43;1.52) | 0.507 | 0.984 |
|  | rs9293503 | 0.97 | (0.5;1.87) | 0.930 | 0.984 |

OR, odds ratio. CI, confidence interval.

Supplementary Table 11	MR-Egger test for directional pleiotropy

| Outcome | Exposure | Intercept | SE | *P*-value |
| --- | --- | --- | --- | --- |
| Overall lung cancer | Self-reported moderate-to-vigorous | 0.022 | 0.020 | 0.290 |
|  | Accelerometer-measured (‘average acceleration’) | -0.011 | 0.028 | 0.713 |
|  | Accelerometer-measured (‘overall activity’) | -0.006 | 0.050 | 0.913 |
| Adenocarcinoma | Self-reported moderate-to-vigorous | -0.017 | 0.031 | 0.593 |
|  | Accelerometer-measured (‘average acceleration’) | -0.008 | 0.042 | 0.855 |
|  | Accelerometer-measured (‘overall activity’) | 0.067 | 0.066 | 0.381 |
| Squamous cell carcinoma | Self-reported moderate-to-vigorous | 0.029 | 0.037 | 0.441 |
|  | Accelerometer-measured (‘average acceleration’) | -0.069 | 0.047 | 0.198 |
|  | Accelerometer-measured (‘overall activity’) | -0.091 | 0.089 | 0.382 |
| Small cell carcinoma | Self-reported moderate-to-vigorous | 0.103 | 0.057 | 0.087 |
|  | Accelerometer-measured (‘average acceleration’) | 0.083 | 0.070 | 0.297 |
|  | Accelerometer-measured (‘overall activity’) | -0.108 | 0.175 | 0.582 |
| Never smoker | Self-reported moderate-to-vigorous | 0.049 | 0.080 | 0.546 |
|  | Accelerometer-measured (‘average acceleration’) | 0.010 | 0.075 | 0.899 |
|  | Accelerometer-measured (‘overall activity’) | 0.079 | 0.140 | 0.614 |
| Ever smoker | Self-reported moderate-to-vigorous | 0.015 | 0.026 | 0.566 |
|  | Accelerometer-measured (‘average acceleration’) | 0.005 | 0.034 | 0.896 |
|  | Accelerometer-measured (‘overall activity’) | 0.034 | 0.065 | 0.633 |

SE, standard error.

1.	Klimentidis YC, Raichlen DA, Bea J, Garcia DO, Wineinger NE, Mandarino LJ*, et al.* Genome-wide association study of habitual physical activity in over 377,000 UK Biobank participants identifies multiple variants including CADM2 and APOE. Int J Obes (Lond) **2018**;42:1161-76

2.	Doherty A, Smith-Byrne K, Ferreira T, Holmes MV, Holmes C, Pulit SL*, et al.* GWAS identifies 14 loci for device-measured physical activity and sleep duration. Nat Commun **2018**;9:5257
